# Enhanced AkaLuc Bioluminescence Imaging for Longitudinal Intravital Monitoring of Minimal Residual Disease in a Murine Model of Triple-negative Breast Cancer

**DOI:** 10.1101/2025.01.18.633734

**Authors:** Silvia Steinbauer, Jamie D. Cowles, Mohammad Ali Sabbaghi, Marle Poppelaars, Azaz Hussain, Marina Wagesreither, Daniela Laimer-Gruber, Jozsef Tovari, Gergely Szakacs, Agnes Csiszar

## Abstract

Triple-negative breast cancer (TNBC) is an aggressive form of cancer with poor prognosis. Beyond the absence of targeted therapies, a major challenge is its high recurrence rate, driven by the outgrowth of residual tumor cells that survive chemotherapy and persist during minimal residual disease (MRD). To monitor the therapy response of TNBC by enhanced intravital imaging, we established a clinically relevant combination chemotherapy protocol for the treatment of mouse mammary tumors engrafted from K14cre;Brca1^F/F^;Trp53^F/F^ (KB1P) organoids engineered to express an mCherry-AkaLuc dual reporter (mCA-KB1P). Reproducible MRD and relapse response patterns with significantly extended relapse-free survivals were achieved with the TAC protocol, consisting of docetaxel, doxorubicin and cyclophosphamide. AkaLuc bioluminescence imaging (AkaBLI) of mCA-KB1P organoids verified the single-cell sensitivity of the system *in vitro,* and showed a detection limit of approximately 1000 cells in the mammary gland of living mice. Unexpectedly, mCA-KB1P organoids elicited an immune response, which necessitated the use of immunodeficient hosts for the longitudinal intravital monitoring of MRD. AkaBLI and an adapted TAC protocol enabled, for the first time, the non-invasive intravital tracking and an estimation of the number of surviving tumor cells in the MRD state following intensive chemotherapy. Engineering KB1P organoids for Histon2B-mCherry reporter expression (HmC-KB1P) enabled the estimation of tumor cell survival also in syngeneic immunocompetent hosts. Flow cytometry and histological analysis revealed that immunocompetent hosts harbored only a few residual cells at MRD, which exhibited a transient loss of epithelial characteristics, whereas immunodeficient hosts had a greater number of surviving cells with a maintained epithelial phenotype. These findings are consistent with the immune system’s role in shaping phenotypic changes influencing survival following chemotherapy Together, the results demonstrate the utility of the AkaBLI system for rare tumor cell tracking and highlight the immune system’s role in triggering adaptive responses to chemotherapy.

## 1. Introduction

Breast cancer, with 2.3 million new cases and over 650,000 deaths in 2022, is the leading cause of cancer-related mortality in women[1]. While well-established diagnostic and therapeutic guidelines exist for estrogen receptor- (ER), progesterone receptor- (PR), and human epidermal growth factor receptor-2- (HER2) positive breast cancers[2], triple-negative breast cancer (TNBC) remains a significant clinical challenge due to the absence of targeted treatment options [3-5]. Treatment of TNBC primarily involves chemotherapy combined with surgery and radiation, while emerging targeted therapies and immunotherapy are under active investigation[5, 6]. The European Society for Medical Oncology (ESMO) Guideline Committee has established standard anthracycline-based regimens for TNBC management. According to their recommendations, preferred approaches include doxorubicin (DOX)- cyclophosphamide (CP)or epirubicin-CP, administered over four cycles spanning 8 to 12 weeks, followed by a taxane for an additional four cycles[7, 8]. The combination of DOX, carboplatin (CPL), and CP is under investigation for its efficacy in neoadjuvant therapy[8].

For neoadjuvant chemotherapy in TNBC, achieving a pathological complete response (pCR) is a reliable surrogate marker for long-term outcomes, including improved response and survival. Despite higher pCR rates, patients with TNBC experience significantly worse overall survival (OS) compared to those with non-TNBC breast tumors[9]. Consequently, disease recurrence remains a critical concern for TNBC patients, with relapse rates typically peaking approximately three years post-diagnosis[10]. Treatment resistance can arise from intrinsically resistant cells, or from the survival of rare residual tumor cells that endure treatment and persist through a prolonged minimal residual disease (MRD) stage[11-13]. While resistant cells continue to proliferate despite cytotoxic therapy, drug-tolerant persister (DTP) cells undergo non-genetic adaptations, entering a dormant or slow-proliferating state[14]. DTP cells display diverse phenotypes, including metabolic changes, epithelial-to-mesenchymal transition (EMT), and activation of various detoxification pathways[15]. Over time, DTPs may transition into fully resistant populations, so early intervention is critical to disrupt this progression[16].

Due to the difficulty in acquiring relevant patient samples, preclinical research on MRD primarily depends on *in vitro* and *ex vivo* systems, mouse models, and patient-derived xenografts[17, 18]. Genetically engineered mouse models (GEMMs) are particularly useful tools as they replicate key aspects of human malignancies, including genetic profiles, interactions with the tumor microenvironment, and therapeutic responses[19]. BRCA1 mutation carriers have a high likelihood of developing TNBC[20]. A prominent model of BRCA1-deficient TNBC is the K14cre;Brca1^F/F^;Trp53^F/F^ (KB1P) mouse model, which has significantly advanced our understanding of therapy resistance mechanisms in BRCA1-mutated breast cancer[21]. Using Cre/loxP technology, this GEMM induces deletions of the BRCA1 and Trp53 genes within keratin 14-positive epithelial cells, leading to the development of spontaneous mammary tumors that closely mimic the histopathological features and therapeutic responses of human BRCA1-mutated cancers[22]. Like TNBC patients, tumor-bearing KB1P mice initially respond to chemotherapy but eventually acquire resistance to multiple treatments[23-25]. Three-dimensional cancer-organoids have also been derived from KB1P tumors[26], enabling *in vitro* culture, genetic manipulation and clinically relevant mechanistic studies on therapy response and resistance[24, 26-27].

Accurate assessment of the residual tumor load after neoadjuvant chemotherapy is a crucial prognostic factor for determining outcome and survival. In mouse models, detecting minimal residual disease requires enhanced resolution and tissue penetration. Recent advancements in bioluminescence-based imaging technologies have facilitated non-invasive preclinical imaging with single-cell resolution[28]. Notably, directed evolution of firefly luciferase led to the development of the AkaLuc system, which generates in vivo emissions that are 100 to 1000 times brighter than those of conventional systems, allowing noninvasive visualization of single cancer cells trapped within the lungs of mice[29]. Here, we apply a combination treatment mimicking neoadjuvant chemotherapy for TNBC patients to follow tumor cells through minimal residual disease and relapse in mice bearing organoid-derived BRCA-deficient mammary tumors using enhanced, non-invasive BLI. In wild-type hosts, analysis of minimal residual disease cells revealed a transient loss of EpCAM positivity. In contrast, immunodeficient hosts exhibited a higher number of MRD cells retaining an epithelial phenotype. Together, these findings demonstrate the utility of the enhanced bioluminescence system for rare tumor cell tracking and highlight the immune system’s role in shaping adaptive responses to chemotherapy.

## 2. Methods

### 2.1 mCA plasmid generation

The bicistronic dual-reporter lentiviral construct was generated using the NEBuilder HIFI DNA Assembly Cloning Kit (#E5520S, New England Biolabs) with the lentiviral vector pRRL-EF1a-WPRE[30] as backbone, linearized with the restriction enzymes BamHI (R3138S, New England Biolabs) and SalI (R3136S, New England Biolabs). Vector pU6-(BbsI)_CBh-Cas9-T2A-mcherry-P2A-Ad4E4orf6[31] was a gift from Ralf Kuehn (Addgene plasmid # 64222; http://n2t.net/addgene:64222; RRID:Addgene_64222) and was used for polymerase chain reaction (PCR) amplification of the mCherry-T2A. Vector pcDNA3 Venus-AkaLuc[29] was provided by the RIKEN BRC through the National BioResource Project of the MEXT, Japan (cat. RDB15781) and was used for PCR amplification of the AkaLuc fragment. Sequence fidelity was confirmed after plasmid assembly by Sanger sequencing.

### 2.2 Cell and organoid culture

HEK293FT (RRID: CVCL_6911) cells were used for lentivirus production. Parental and engineered 4T1[32] and MCF7 (ATCC HTB-22) cells were used for *in vitro* and *in vivo* titration assays. Cell lines were maintained in Dulbecco’s modified Eagle medium (DMEM; HEK293FT and 4T1) or RPMI (MCF7) supplemented with 10% fetal bovine serum (FBS), and 1% penicillin/streptomycin. During transfection, cells were grown without antibiotics. Parental and engineered KB1P breast cancer organoids were cultured as described[26]. Cells and organoids were maintained at 37°C with 5% CO_2_.

### 2.3 Lentiviral production in HEK293FT cells

The mCA constructs were used to produce lentivirus in HEK293FT cells using the standard CaCl_2_ transfection method. Briefly, 6×10^6^ HEK293FT cells were seeded into T75 culture flasks. After 24 hours, the second-generation lentiviral packaging vectors psPAX.2 and PMD2.G and the plasmid of interest (mCA-plasmid, pLenti.PGK.H2B-chFP.W[33], or pHIV-iRFP720-E2A-Luc[34]) were slowly mixed with sodium-phosphate. The generated DNA-calcium phosphate precipitate was directly pipetted onto the cells, which were cultivated in antibiotic-free medium for a few hours. 24 hours post transfection, fresh medium was added, and the cells were incubated for an additional 48 hours. Finally, the supernatant containing the viral particles was collected and filtered with a 0.45 µm low protein binding filter. For viral concentration, the filtered supernatant was ultra-centrifuged at 25,000 rpm for 2 hours at 4°C, and the pelleted virus was resuspended in 1/100 of the original volume. Viral titers were determined using the abm qPCR Lentivirus Titration Kit (abm, #LV900) according to the instructions. psPAX2 and PMD2.G were a gift from Didier Trono (Addgene plasmid # 12260; http://n2t.net/addgene:12260; RRID:Addgene_12260 and Addgene plasmid # 12259; http://n2t.net/addgene:12259; RRID:Addgene_12259, respectively). pLenti.PGK.H2B-chFP.W[33] was a gift from Rusty Lansford (Addgene plasmid # 51007; http://n2t.net/addgene:51007; RRID:Addgene_51007). pHIV-iRFP720-E2A-Luc[34] was a gift from Antonius Plagge (Addgene plasmid # 104587; http://n2t.net/addgene:104587; RRID:Addgene_104587).

### 2.4 Lentiviral transduction of cell and organoid lines

Concentrated mCA viral particles at a multiplicity of infection (MOI) of 25 were added to 9×10^4^ MCF-7 cells in a final volume of 1 mL cultivation media in one well of a 6-well plate. After 48 hours incubation at 37°C with 5% CO_2_, cells were expanded and further cultivated according to standard procedures.

Lentiviral transduction of KB1P organoids was performed according to Duarte et al.[22] with the following modifications: concentrated viral particles were mixed to 5×10^5^ (mCA) or 1.2×10^5^ (HmC, or FLuc) single cells of KB1P organoids at a MOI 25 in a final volume of 500 µL ENR in one well of an ultralow adherent 24-well plate (Greiner Bio-ONE CELLSTAR, #391-3375), followed by spinfection for 1 h at 600 g, room temperature and incubation for 6 hours (mCA) and 48 hours (HmC, or FLuc), respectively, at 37°C with 5% CO2. Thereafter, KB1P cells were seeded in a density of 3×10^4^ cells/40 µL matrix in BME:ENR (1:1), and further cultivated according the original protocol.

Transduction efficiency of mCA, HmC, or FLuc was evaluated by flow cytometry. To enrich for transduced cells, Fluorescence Activated Cell Sorting (FACS) was applied, followed by flow cytometric validation of enrichment. Finally, reporter expression was evaluated by microscopy.

### 2.5 Mice and mammary fat pad transplantation

Mice were kept in the animal facility of the Medical University of Vienna in accordance with institutional policies and federal guidelines. Animal experiments were approved by the Animal Experimental Ethics Committee of the Medical University of Vienna and the Austrian Federal Ministry of Science and Research (animal license number: BMBWF 2020-0.121.294 and BMBWF 2024-0.518.920). Organoids were transplanted as described earlier[26]. Briefly, organoids were used at a size corresponding to an average of 150–200 cells per organoid structure. Organoid suspensions containing a total of 5 × 10^4^ cells in 30 µl of organoid media/BME mixture (1:1) were injected orthotopically into the fourth right mammary fat pad of 7-week-old wild-type FVB/N (KB1P, HmC-KB1P, mCA-KB1P), NMRI nude (mCA-KB1P), and NSG (mCA-KB1P) mice (Charles River Laboratories, Germany) under anesthesia (100 mg/kg ketamine and 5 mg/kg xylazine) after a small surgical incision. The tumor size was monitored at least 3 times per week by caliper measurements. Tumor volumes were calculated using the formula V = 0.5 x length x width^2^. Animals were sacrificed at the experimental endpoint (200-300 mm^3^ for drug-naïve, at MRD, or 200-2000 mm^3^ for relapse samples). For engraftment experiments mice were sacrificed before the tumor volume reached 2000 mm^3^.

### 2.6 Chemotherapeutics and treatment of tumor-bearing mice

DOX (Adriblastin; 2 mg/ml), pegylated liposomal DOX (Doxil) (Caelyx®; 2 mg/ml), Docetaxel (DTX; Accord Healthcare; 20 mg/ml), and CP (Endoxan; 20 mg/ml) were obtained directly from the manufacturers in clinically used formulations and were further diluted in saline before use to a final injection volume of 200 µL.

The triple-combination therapy TAC was performed in FVB/N mice as described earlier for other tumor models[35] with modifications. Briefly, the drugs were used in following doses: docetaxel 25 mg/kg, DOX 5 mg/kg and CP 120 mg/kg. First, CP was administered by intraperitoneal (i.p.) injection, which was followed by an intravenous (i.v.) injection of a freshly prepared mix of DTX and DOX. Therapy was started either at a tumor volume of 200 ± 50 mm^3^, including two treatment cycles separated by 21-days (TAC(2x,q21)); or at 100 ± 25 mm^3^, with two cycles separated by 5 days (TAC(2x,q5)). Nude mice were treated with the same dose. In the case of NSG mice, doses were reduced as follows: DTX 7.5 mg/kg, DOX 1.5 mg/kg and CP 120 mg/kg. For both strains therapy was administered at a tumor volume of 100 ± 25 mm^3^, in two treatment cycles, separated by a 5-day-interval.

### 2.7 Bioluminescence imaging

For *in vitro* evaluation of the AkaBLI sensitivity in the mCA-KB1P organoids, an *in vitro* titration assay was combined with BLI. Parental KB1P, mCA-KB1P and FLuc-KB1P organoids and parental MCF7, parental 4T1, mCA-MCF7 and FLuc-4T1 cells were trypsinized to single cells and serial dilutions of 5, 10, 50, 100, 250, 500 and 1000 cells were seeded in PBS/0.5% BSA in triplicates into a black 96-well imaging plate with a glass bottom (Eppendorf, #0030741030). AkaLumine (#013-23693, FUJIFILM Wako Chemicals Europe GmbH) substrate solution [500 µM] or D-Luciferin (#7902, BioVision) substrate solution [99 mM], was added to the cells in a final concentration of 250 µM. Bioluminescence measurements were performed after 15 min of substrate incubation as described previously[29]. The following settings were used for the IVIS® Spectrum System (Perkin Elmer): open for total bioluminescence, exposure time: 40 secs (MCF-7, KB1P, mCA-MCF-7, mCA-KB1P), 60 secs (FLuc-KB1P), and 300 secs (4T1, FLuc-4T1), binning: 4 (medium), FOV: C, f/stop: 1, temp at measurement: 20-25°C.

To evaluate drug-induced intensity changes of AkaBLI, in vitro 2D repopulation assay was performed in parental and mCA-MCF7 cells as described[15] and cells were harvested by trypsinization for BLI measurements at day 1 (drug naïve), 22, 29 (DTP) and 60 (Repop). AkaBLI was measured in a plate reader (Tecan) after seeding 10,000 cells in PBS/0.5% BSA in triplicates into a black 96-well imaging plate with glass bottom (Corning) and immediate addition of AkaLumine in a final concentration of 250 µM with following settings: central wavelength of 648 nm with 105 nm bandwidth, integration time of 3000 ms and kinetic mode with four measurement cycles in two minutes intervals. Technical triplicates were averaged and a minimum of three independent biological replicates were measured.

*In vivo* titration analyses were performed to test the *in vivo* sensitivity of the bioluminescence of the mCA-KB1P organoids. Engineered KB1P organoids and engineered 4T1 cells were trypsinized to single cells and serial dilutions of 1×10^2^, 5×10^2^, 1×10^3^, 5×10^3^, 1×10^4^, 5×10^4^, 1×10^5^, 2.5×10^5^, 5×10^5^ organoid cells were prepared in PBS on ice. A volume of 30 µL of these organoid-cell solutions was injected into the fourth left and right mammary fat pad of anaesthetized female BalbC, nude, or NSG mice. 17 min after i.p. injection of 50 mg/kg AkaLumine-Hydrochloride (“TokeOni”, #HY-112641A, THP and #808350, Sigma Aldrich) substrate (30 mM in ddH2O) or 150 mg/kg D-Luciferin (#7902, BioVision) substrate [99 mM] solution, bioluminescence measurements were performed at the IVIS® Spectrum System (Perkin Elmer) with the following settings: open for total bioluminescence, exposure time: 1 sec, binning: 4 (medium), FOV: D, f/stop: 1, temp at measurement: 20-25°C.

For *in vivo* detection of transplanted, engrafted, treated, and relapsed mCA-KB1P tumors, BLI was performed the day after transplantation, the day of drug treatment (75-150 mm^3^ tumor size), after chemotherapeutic treatment at MRD (< 5 mm^3^ tumor size), and when the tumors relapsed (100-600 mm^3^ tumor size). BLI analysis was performed in Living Image (Perkin Elmer) software.

### 2.8 Tumor dissociation and flow cytometry

Tumor-bearing mice were sacrificed before the initiation of the treatment (Drug-Naïve, DN), at the MRD stage, or at relapse (REL). While DN and REL tumors could be dissected as well-defined tissue masses, in MRD samples a comparable, 1,5 cm long central piece of the mammary fat pad (site of injection at transplantation) around the inguinal lymph node was cut out, the adjacent normal tissue providing carrier cells for the preparation. Suspensions containing a high rate of viable cells were prepared as described[36], with modifications. Briefly, excised tissue was cut into small pieces using scalpel blades, tissue fragments were shaken in Digestion Media (RPMI supplemented with 250 µg/ml DNase I and 1 mg/ml Collagenase IV) for 40 minutes at 37 °C. Digests were filtered through 70 µm strainers, washed with DNase I (100 µg/ml) containing washing buffer (PBS with 2% FCS), and centrifuged for 5 minutes (4 °C, 300g). Following lysis of red blood cells (RBC lysis buffer, # B420301, Biozym) the pellet was washed twice in 10 ml Wash Buffer, centrifuged for 5 minutes (4 °C, 300g). For flow cytometry, the resulting pellet was pre-blocked with anti-CD16/32 antibody (#156603, BioLegend) and stained with the following fluorophore conjugated primary antibodies for 30 min at 4°C: CD31 (PE, #561073, BioLegend), CD45 (BV421, #103134, BioLegend) and epithelial cell adhesion molecule (EpCAM) (Fluorescein isothiocyanate [FITC], #118207, BioLegend), followed by Zombie Aqua viability dye (#77143, BioLegend) staining according to the manufacturer’s recommendation to exclude dead cells. Samples were analyzed by a Fortessa X20 (BD Bioscience) cytometer or by a FACSMelody cell sorter (BD Biosciences); data were analyzed using the FlowJo v10 software (BD Life Sciences).

### 2.9 Immunohistochemistry (IHC)

Immunohistochemistry was performed according to Malik et al.[37]. In brief, following treatment with 3% H_2_O_2_ (Sigma Aldrich), 4-µm thick sections of mouse tumor tissues were repeatedly washed with 0.1% Tween 20 in PBS (PBS-T) and incubated in 5% horse blocking solution for 60 min in a wet chamber. Primary antibodies anti-E-cadherin (1:1000, #3195, Cell Signaling Technologies), anti-Cytokeratin 14 (1:500, #905303, BioLegend), anti-mCherry (1:250, #600-401-379S, ThermoFisher), anti-Ki67 (1:3000, #ab15580, Abcam) and anti-activated Caspase 3 (1:300, #9661S, Cell Signaling Technologies) were diluted in blocking buffer (2% BSA, 5% horse serum in PBS-T) and incubated overnight at 4°C. Slides were washed and incubated with horseradish peroxidase (HRP)-conjugated secondary antibodies (1:2, #8114, Cell Signaling Technologies) for 60 min at room temperature, followed with 3,3′-diaminobenzidine (DAB substrate (DAKO)) for detection according to the manufacturer’s instruction.

### 2.10 Statistical analysis

Statistical analysis was done with GraphPad Prism 10.04. A sample size estimate was not statistically performed. To compare one parameter between two groups unpaired and paired two-tailed Student’s t-test were used, across multiple groups one-way or two-way analysis of variance (ANOVA) with Tukey multiple comparison test was applied to determine statistical significance. OS and RFS were calculated using log-rank statistics. p-values of lower than 0.05 were considered statistically significant (*plJ<lJ0.05, **plJ<lJ0.01, ***plJ<lJ0.001, ****p<0.0001). Error bars are represented as standard error of mean (SEM). The strength of relationship between two variables was calculated by Pearson correlation analysis, linear regression curves calculated and presented with 99% confidence intervals.

## 3. Results

### 3.1 Establishment of a clinically relevant chemotherapy treatment protocol achieving sustained MRD and predictable relapse of organoid-derived mammary tumors

To model the different phases of clinical therapy response in breast cancer, mouse mammary tumors, derived from KB1P organoids previously established from a BRCA1/p53 deficient murine breast tumor[26], were treated with chemotherapy. Orthotopic transplantations of KB1P organoid structures into the 4^th^ mammary fat pad of syngeneic FVB/N female mice showed reproducible growth kinetics (**Figure 1A**) that matched the characteristics of the model originally published by Duarte et al.[26]. To establish treatment schedules resulting in reproducible response patterns, including a stable MRD phase followed by relapse, organoid-derived tumors were treated with DOX according to treatment protocols established previously for the KB1P model[24]. Though DOX therapy increased the median OS of KB1P tumor-bearing mice (**Figure 1B**), it did not lead to a significant decline of the tumor size (**Figure 1C**). Our earlier work demonstrated that treatment with the liposomal formulation of DOX (pegylated liposomal DOX; Doxil Caelyx®) is effective even against DOX refractory KB1P tumors[24]. Accordingly, treatment with Doxil led to a significant increase of the median OS of tumor-bearing mice compared to DOX treated animals (**Figure 1B**). Albeit several mice treated with Doxil showed complete response, the onset and duration of the MRD phase varied (**Figure 1D**). To ensure reproducible and higly similar response patterns, we adapted a clinically used combination treatment, consisting of DOX, DTX, and CP (TAC protocol), previously described for the MMTV-PyMT breast cancer model [35]. Indeed, two TAC treatments separated by 21 days proved sufficient to induce a stable MRD phase in every tumor bearing mouse, resulting in significantly increased OS and a median RFS of 96 days (**Figure 1E,F; TAC(2x,q21)**). However, the onset of the relapse remained unpredictable, with some tumors relapsing before the administration of the second TAC cycle (**Figure 1G**). To circumvent early relapse and improve synchronization of therapy response, the TAC protocol was modified by administering the second treatment cycle earlier, after 5 days (TAC(2x,q5)). While the higher initial cumulative drug dose of this treatment protocol resulted in decreased OS and RFS compared to the TAC(2x,q21) protocol (**Figure 1E,F**), this modification increased the short-term efficacy of the treatment without imposing additional toxicity. Importantly, the TAC(2x,q5) protocol resulted in the reproducible induction of a stable MRD phase with a predictable length of 30-40 days after treatment start in 70% of the animals (**Figure 1F,H**). Taken together, the establishment of the TAC(2x,q5) combination chemotherapy protocol enabled uniform response patterns of murine KB1P organoid-derived mammary tumors, characterized by the reproducible induction of a stable MRD phase with predictable duration before tumor relapse.

**Figure 1.**
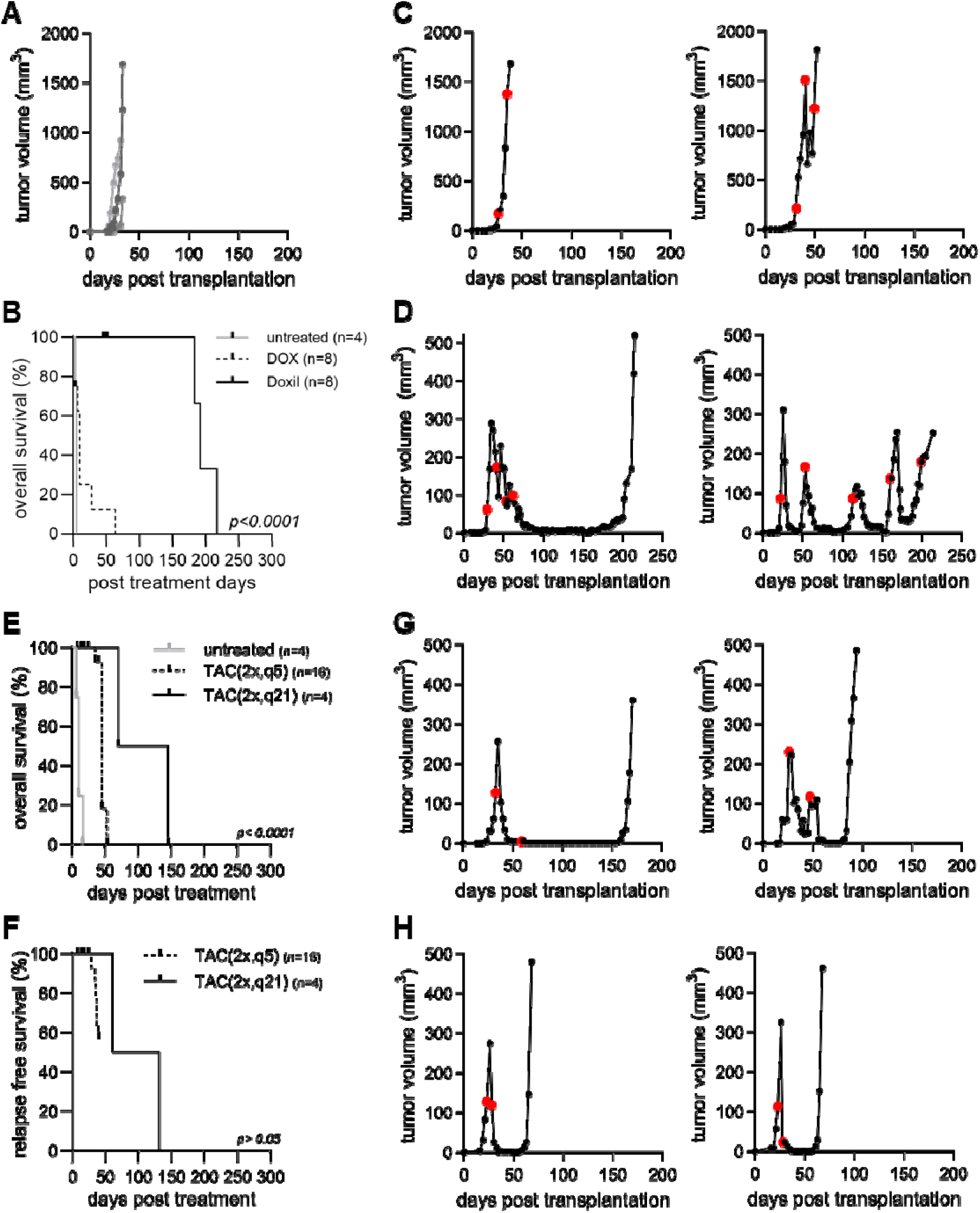
Establishment of a clinically relevant chemotherapy protocol with a stable MRD in the syngeneic KB1P organoid-derived tumor model. **(A)** Representative tumor growth curves of untreated FVB/N mice after orthotopic transplantation with KB1P tumor organoids. **(B)** Overall survival (OS) of untreated (grey), DOX (dashed line) and Doxil (solid line) treated tumor-bearing FVB/N mice. **(C,D)** Representative tumor growth curves of **(C)** DOX and **(D)** Doxil-treated tumor-bearing FVB/N mice. **(E)** Overall survival (OS) of untreated (grey) and TAC treated tumor-bearing FVB/N mice, second treatment separated by 21 (2x,q21; solid line) or 5 days (2x,q5; dashed line). **(F)** Relapse-free survival (RFS) of KB1P-transplanted FVB/N mice, treated with the two different TAC protocols. **(G,H)** Representative tumor growth and drug response curves of KB1P-transplanted FVB/N mice, treated with **(G)** TAC(2x,q21) or the **(H)** TAC(2x,q5) protocols. C,D,G,H red circles on the tumor growth curves mark treatment days with the respective drugs.

### 3.2 KB1P organoids engineered to express an mCherry-AkaLuc dual reporter confirm the high detection sensitivity of the AkaLuc/AkaLumine system *in vitro*

To detect the few surviving cells in the non-palpable MRD stage, we aimed to establish high-sensitivity bioluminescence as an intravital imaging tool. We generated a bicistronic lentiviral dual reporter construct expressing AkaLuc, a luciferase with single-cell detection sensitivity in living animals[29] and mCherry as a fluorescent marker (**Figure 2A**). Fluorescence microscopy and fluorescence-activated cell sorting (FACS) analysis confirmed the stable expression of the reporter construct in transduced KB1P organoids (mCA-KB1P) after two rounds of enrichment by FACS (**Figure 2B,C**). *In vitro* luciferase assays validated the AkaLuc expression and confirmed a 100-fold increased *in vitro* sensitivity of the AkaLuc/AkaLumine system compared to Firefly luciferase (FLuc)/Luciferin[38] (**Figure 2D**). The lowest tested cell number of 5 cells could be detected with the AkaLuc/AkaLumine system compared to a detection limit of 500 cells for KB1P organoids engineered with FLuc. This difference in the detection sensitivity remained consistent with increasing cell numbers and appeared independent of the cell line (**Figure 2E).**

**Figure 2.**
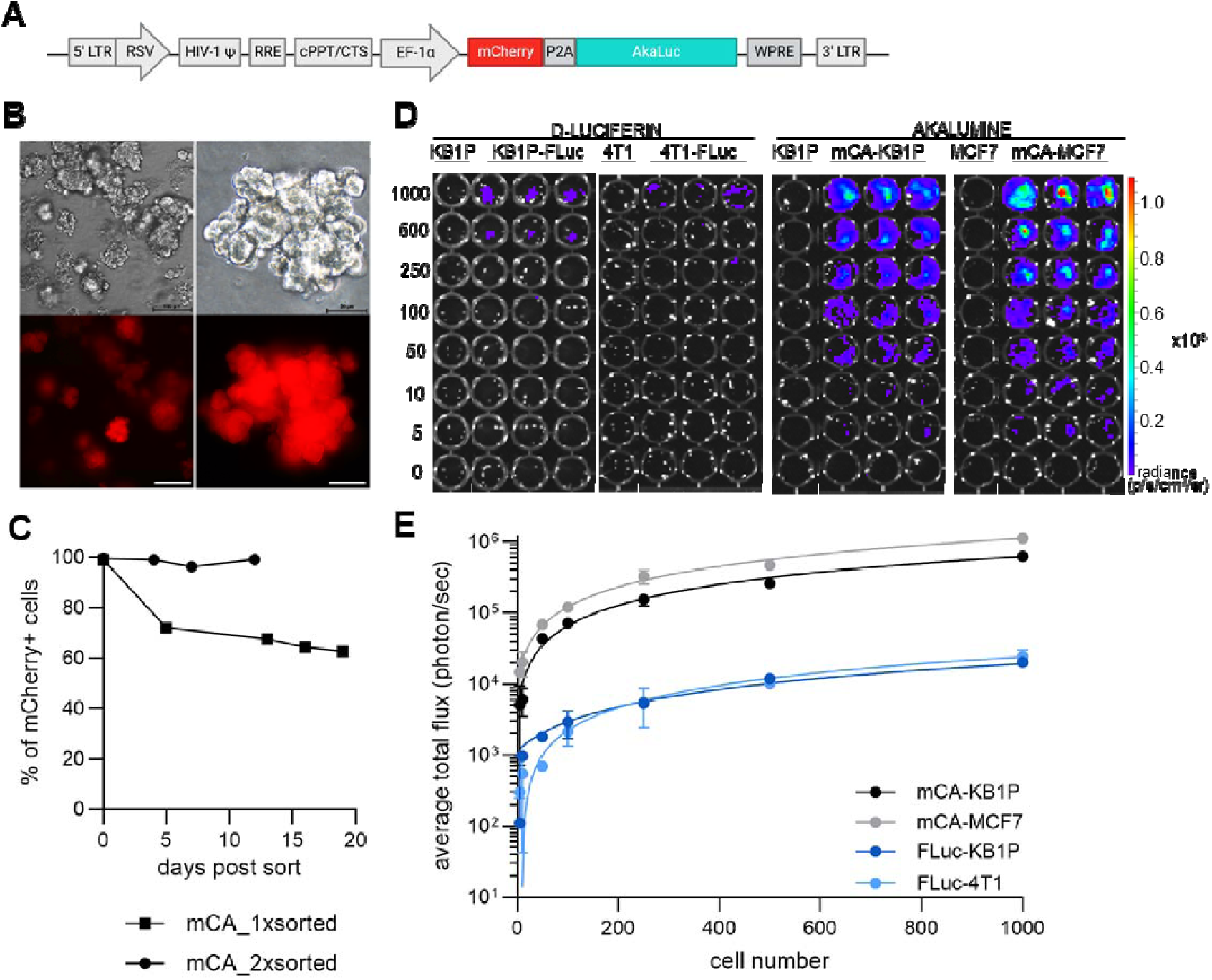
Generation and in vitro detection sensitivity of the dual mCherry-AkaLuc reporter system in KB1P organoids. **(A)** Schematic representation of the mCherry-AkaLuc lentiviral reporter construct. **(B)** Representative bright field and fluorescence microscopy images of KB1P organoids stably transduced with the mCherry-AkaLuc reporter (mCA-KB1P). Scale bar, 100 µm (left panel) and 20 µm (right panel). **(C)** Longitudinal FACS-assisted stability testing of the reporter in mCA-KB1P organoids. **(D)** Representative image of bioluminescence intensity (depicted by the color scale) with increasing cell numbers (indicated by the numerical values on the left) for KB1P organoids and 4T1 cells with Fluc, and KB1P organoids and MCF7 cells with AkaLuc expression. Wells containing parental organoids/cell lines served as negative controls. **(E)** Quantification of in vitro bioluminescence as total flux/cell [p/s/cell] as a function of plated cell number.

### 3.3 Adapted TAC chemotherapy protocol for immunodeficient hosts enables the study of rare surviving MRD cell populations despite the immunogenicity of AkaLuc

The high detection sensitivity of the AkaLuc reporter, validated in the *in vitro* KB1P organoid model, paved the way for monitoring the few tumor cells that survive chemotherapy during the non-palpable MRD stage in living animals using BLI. Unexpectedly, mCA-KB1P organoids failed to engraft in syngeneic FVB/N mice (**Figure 3A**). To determine whether this was due to a loss of tumorigenic capacity of engineered organoids or a potential immunogenic reaction of the host against the AkaLuc reporter, we transplanted parental KB1P and mCA-KB1P organoids into immunocompromised nude and immunodeficient NSG mice (**Figure 3B,C).** mCA-KB1P organoids efficiently formed tumors in both immunodeficient strains, demonstrating their preserved tumorigenicity (**Figure 3D**). These observations strongly suggest an immunogenic characteristic of the AkaLuc protein, which has not been reported previously. The moderate engraftment rate in nude mice (81%) compared to NSG mice (100%), along with the significantly delayed tumor outgrowth in both strains relative to the parental line in immunocompetent FVB/N hosts (**Figure 3D-E**), provides further indication of a complex immune surveillance mechanism targeting AkaLuc, with T-cells playing an important, but not exclusive role. Together, these transplantation experiments highlight the immunogenic characteristic of the AkaLuc protein, rendering it unsuitable for use in immunocompetent transplantation hosts.

**Figure 3.**
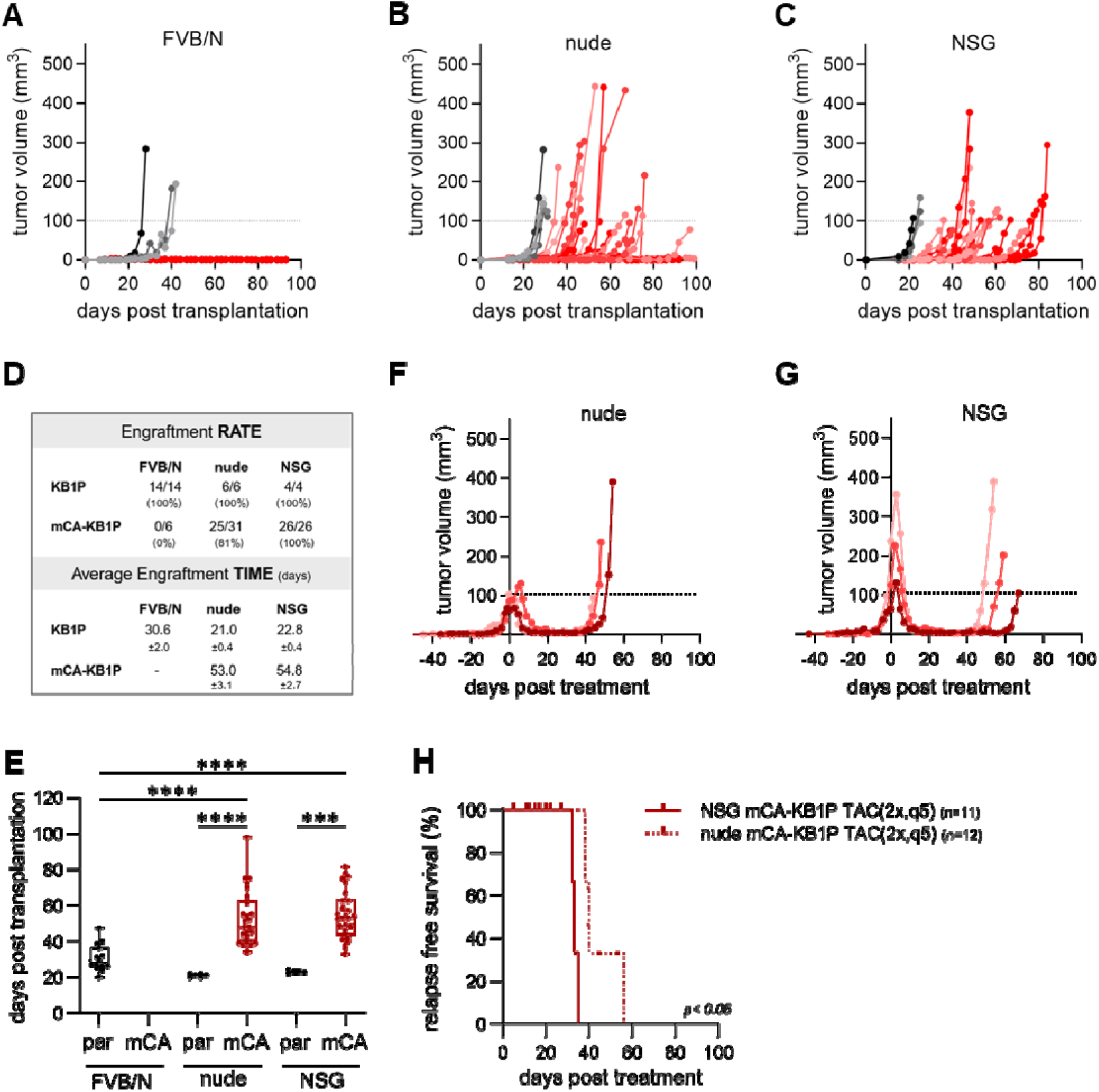
Immunodeficient mice are suitable hosts for mCA-KB1P tumor engraftment and the modeling of MRD following adapted TAC chemotherapy. **(A-C)** Tumor growth curves obtained in **(A)** FVB/N, **(B)** nude and **(C)** NSG mice transplanted with parental KB1P (grey shaded lines) or mCA-KB1P (red shaded lines) organoids. **(D)** Statistics of the engraftment rate (%) and time (days) of the transplants. **(E)** Median engraftment time (days needed after transplantation to reach treatment size). Statistical analysis was performed by One-way ANOVA. Significant differences are marked with asterisks **** p<0.0001; *** p<0.001. “par” and “mCA” mark tumor transplants with KB1P parental and mCA-KB1P organoids, respectively. **(F,G)** Tumor growth and drug response of **(F)** nude and **(G)** NSG mice transplanted with mCA-KB1P organoids and treated with the TAC(2x,q5) protocol **(H)** Relapse-free survival after therapy start of nude and NSG mice transplanted with mCA-KB1P organoids.

To induce the MRD phase in immunodeficient mice, we evaluated the efficacy of the established TAC(2x,q5) protocol in nude and NSG mice. To this end, we first optimized the drug dosing, using the strain-specific maximum tolerated dose (MTD) and existing treatment regimens of the individual components as guidelines [39-42]. Nude mice showed similar drug tolerability, allowing treatment with the same drug dosing as in syngeneic FVB/N hosts. mCA-KB1P tumors in nude mice showed a complete response, characterized by a stable MRD phase, followed by relapse (**Figure 3G**), similar to the therapy response observed with parental tumors in syngeneic FVB/N hosts (**Figure 1H**). NSG mice tolerated a substantially lower dosing of DOX and DTX. Nevertheless, the NSG-adapted TAC(2x,q5) protocol recapitulated the treatment response of KB1P tumors observed in the other strains. All tumor-bearing NSG mice showed a stable MRD of reproducible length (20±5 days) (**Figure 3F**), albeit median RFS was significantly shorter than in nude mice (33 vs. 44 days) (**Figure 3H**). In summary, these data demonstrate the robustness of the mCA-KB1P tumor transplantation model combined with a TAC chemotherapy in immunodeficient hosts. They also highlight the significant role of immune components beyond T cells, including B cells, natural killer (NK) cells, and elements of the complement system, in tumor surveillance during chemotherapy. Importantly, NSG mice did not exhibit severe systemic toxic side effects, highlighting the applicability of the newly established, NSG-adapted TAC protocol in pre-clinical tumor therapy models. Furthermore, this protocol enabled the generation and monitoring of rare surviving tumor cells during the MRD phase in the mCA-KB1P organoid model in NSG mice using AkaBLI.

### 3.4 Enhanced AkaLuc bioluminescence imaging of surviving tumor cells during minimal residual disease

The AkaLuc reporter has been described as providing single-cell sensitivity for bioluminescent detection in living animals[29]. The *in vivo* transplantation assay determined a detection limit of approximately 1,000 cells (**Figure 4A,B**), which was 10 times lower than the 10,000 cell detection limit for KB1P organoids engineered with FLuc, and 50 times lower compared to transplanted 4T1 cells engineered with FLuc (**Figure 4A,B**). Thus, while the *in vivo* sensitivity appeared more moderate than previously described, with potential differences between engineered cell lines and/or host strains, it was still superior by at least one order of magnitude compared to the conventional FLuc system. A limitation of the system is the hepatic background signal of the AkaLumine substrate, as described in [43-46], which was uniformly observed across different mouse strains (**Supplementary Figure S1**). However, this issue did not affect the present study, as the 4^th^ mammary fat pad does not overlap with the hepatic region.

**Figure 4.**
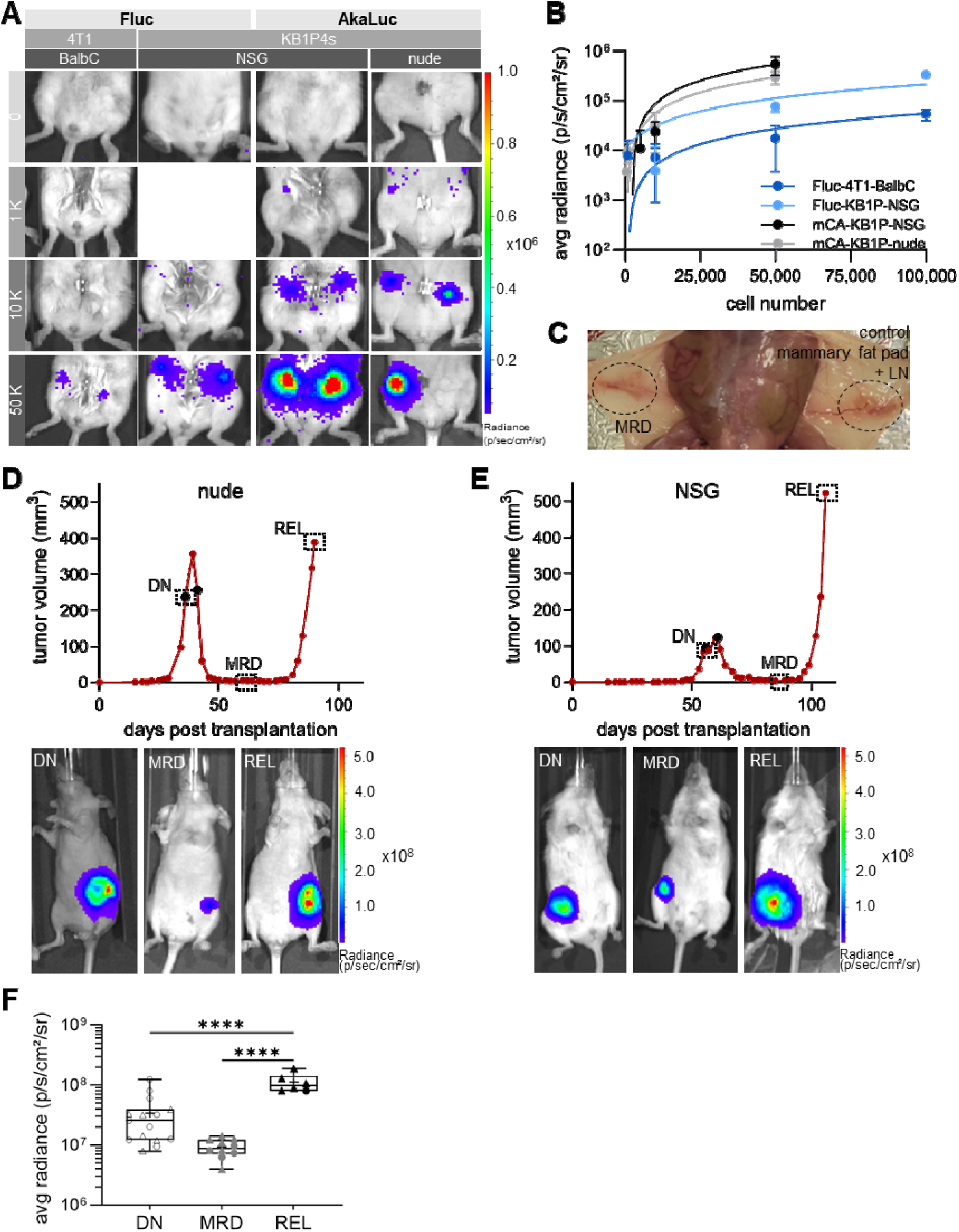
AkaLuc bioluminescence imaging of surviving tumor cells during minimal residual disease. **(A)** Representative bioluminescence images of mice transplanted with increasing numbers of Fluc or AkaLuc expressing tumor cell or organoid lines. **(B)** Dose-titration of Fluc- and AkaLuc-derived bioluminescence in various cell- and organoid lines across different mouse strains, represented as mean radiance relative to transplanted cell numbers. **(C)** Representative macroscopic image of the right mammary fat pad of a nude mouse in the non-palpable MRD stage after mCA-KB1P organoid transplantation. The contralateral organ is shown as non-transplanted control. **(D,E)** Representative growth curves and drug response of mCA-KB1P organoid-derived tumors of a **(D)** nude and **(E)** NSG mouse treated with the TAC(2x,q5) treatment protocol. Treatment days are indicated by black dots. Representative images of longitudinal BLI at drug naïve (DN), MRD, and relapse (REL) stages are shown under the corresponding tumor curves, framed data points mark the respective timepoints of BLI. **(F)** Quantification of BLI signal intensity as mean radiance of mCA-KB1P tumor-bearing nude and NSG mice at DN, MRD and REL stages (nude n=1-9, NSG n=5-6 per time point). One-way ANOVA was used for statistical analysis for combined values of nude and NSG mice. Circles and triangles mark nude and NSG data points, respectively. Significant differences are marked with asterisks, (**** p<0.0001).

The high sensitivity of the AkaLuc-AkaLumine system allowed the robust detection of the 50,000 mCA-KB1P organoids transplanted at the start of the experiment (**Supplementary Figure S2A**). In addition, it provided further experimental support for the eradication of these cells in immunocompetent FVB/N hosts (**Supplementary Figure S2A-C**). Interestingly, it also revealed the long-term survival of transplanted tumor cells in those nude mice, which did not show tumor outgrowth even after 110 days (∼20% of transplants) (**Supplementary Figure S2A**), with the intensity of the initial BLI signal maintained (**Supplementary Figure S2B,C**). This indicates tumor dormancy, as confirmed by IHC analysis of proliferation and apoptosis markers (**Supplementary Figure S2D**), likely due to residual immune surveillance.

Having established transplantation and treatment protocols for studying a stable MRD phase in immunodeficient mice bearing mCA-KB1P expressing tumors, we aimed to detect and quantify surviving tumor cells in MRD by intravital BLI. To monitor BLI signal intensity changes over chemotherapy response, mice transplanted with mCA-KB1P organoids were imaged (i) as soon as tumors reached treatment size (DN stage), (ii) at stable MRD, and (iii) at tumor relapse (REL). Importantly, the MRD stage showed a prolonged period during which tumors were non-palpable (**Figure 4C**), yet yielded detectable BLI signals in all mice. **Figure 4D** and **E** show a representative BLI image series aligned with the TAC(2x,q5) treatment response of mCA-KB1P tumors in longitudinally monitored nude and NSG mice, respectively.

BLI signal intensity and calculated tumor cell numbers based on the titration experiments shown in **Figure 4A**, correlated with tumor volumes for tumors ranging from 0 and 150 mm³ in size (r=0.4906, p<0.01 and r=0.5966, p<0.001, respectively) (**Supplementary Figure S3A,B**), most likely values of bigger tumors being influenced by necrotic and cystic cores. Thus, BLI data from mice with higher tumor volumes were excluded from statistical analysis. (**Supplementary Figure S3C**). Analyses of the 0-150 mm^3^ tumor cohort showed distinct BLI intensities for the three different tumor stages, ranging from an average radiance of 3×10^7^ in the DN state, to approximately 1×10^7^ in the MRD state, and 1×10^8^ in the REL state (**Figure 4F)**. This distinction was also observed in the corresponding tumor cell numbers calculated from the titration experiments (**Supplementary Figure S3D**).

Average radiance levels showed no substantial strain-specific differences, but significantly lower BLI intensities were observed in the DN compared to the REL stages (**Figure 4F**), despite the highly similar tumor volumes of these two stages (**Supplementary Figure S3C**). While an up to 25-fold reduction of the tumor volume in the MRD phase was associated with only a 5-fold reduction in BLI signal intensity, the 25-fold increase in tumor volume in the REL phase corresponded to a significantly higher, up to 10-fold, increase in BLI signal intensity (**Figure 4F**). These results can be explained by assuming stage-dependent differences in bioluminescence signal generation. Indeed, DTP cells generated in the *in vitro* DTP repopulation assay [15] using mCA-MCF7 cells and DOX treatment exhibited a significant, 4-fold lower bioluminescence signal intensity at DN stage compared to DTP cells (**Supplementary Figure S3E**), supporting the notion that drug-induced phenotypic changes increase the efficiency of bioluminescent signal generation. Taken together, enhanced BLI during the MRD phase revealed a high number of surviving tumor cells in immunodeficient mice after chemotherapy, even if the accuracy of these estimates requires further investigation.

### 3.5 HmC-KB1P organoids engraft in syngeneic immunoproficient FVB/N hosts with maintained characteristics of drug response and stable MRD

To allow characterization of tumor cells along treatment with chemotherapy in immunocompetent hosts, H2B-mCherry (HmC) was introduced as a fluorescent reporter gene into KB1P organoids by lentiviral engineering (HmC-KB1P). Orthotopic transplantation of HmC-KB1P organoids into syngeneic FVB/N hosts showed comparable engraftment rates (**Figure 5A – insert**) and growth kinetics to the parental KB1P model (**Figure 5A**). Similarly, treatment of tumors originating from HmC-KB1P organoids with the TAC(2x,q21) and (2x,q5) protocols showed similar OS and RFS rates (**Figure 5B,C**) with comparable drug responses to parental KB1P organoids (**Figure 5D,E**). The therapy response was also comparable to that of the mCA-KB1P organoids in nude and NSG mice (**Figure 3G and Supplementary Table S1**) thereby establishing a system for comparing MRD in immunocompetent and immunocompromised hosts.

**Figure 5.**
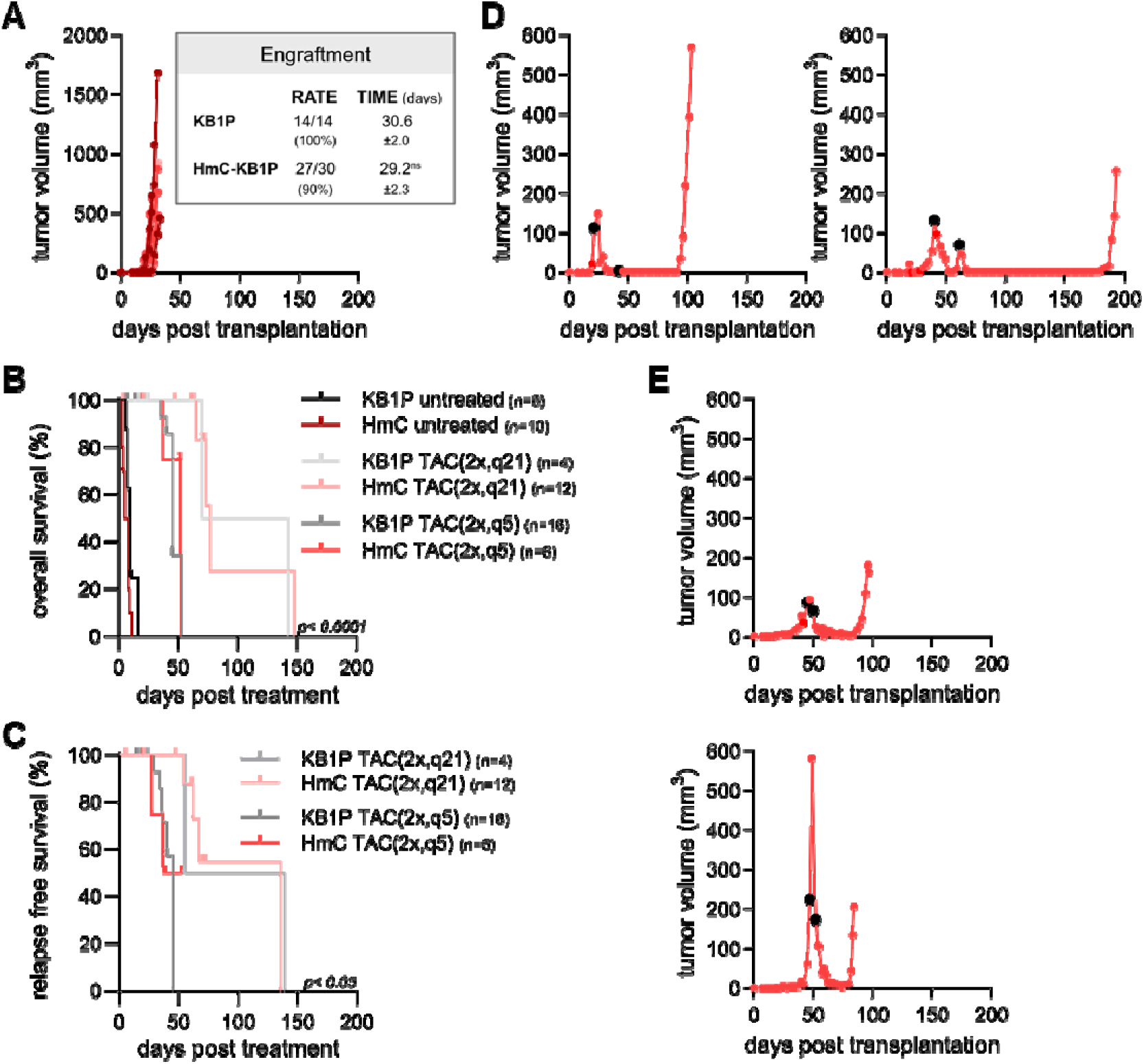
KB1P organoid-derived tumors engineered for HmC reporter expression engraft in syngeneic immunoproficient FVB/N hosts with maintained characteristics of drug response and stable MRD. **(A)** Representative growth curves of orthotopically transplanted HmC-KB1P organoid-derived tumors in female FVB/N mice. Insert shows statistics of the engraftment rate and time of the transplants. Unpaired t-test was used for statistical analysis to compare differences between groups. ns, non-significant. **(B)** OS of untreated (dark lines) and 2x,q21 (light lines) or 2x,q5 (faded lines) TAC protocol treated mice. **(C)** RFS of (HmC)-KB1P-transplanted mice, treated with the two different TAC protocols. **(D,E)** Representative tumor growth and drug response curves of HmC-KB1P-transplanted mice, treated with **(D)** TAC(2x,q21) or **(E)** TAC(2x,q5) protocols. Black dots on the tumor growth curves mark the days of TAC treatment.

### 3.6 The MRD stage is linked to lower adaptive selection pressure in immunocompromised mice

To measure the overall abundance of the surviving tumor cells in the non-palpable MRD tumor stage, six HmC-KB1P and six mCA-KB1P tumors were harvested and pooled at the MRD stage from FVB/N and nude or NSG hosts, respectively. Flow cytometric analysis was then performed using mCherry expression for cancer cell identification (**Figure 6A**), along with the exclusion of immune and endothelial cells to account for differences in the tumor microenvironment (**Supplementary Figure S4A,B**). This analysis revealed that there were fewer tumor cells surviving in FVB/N hosts than in immunodeficient hosts during the MRD phase (0,47% vs. 14,5% of the CD45-,CD31-double-negative live cell population, respectively). This suggests that in immunocompetent mice, only a very small number of cells survive the treatment, serving as a reservoir for replenishing the relapsing tumor. In contrast, as also indicated by the BLI measurements, significantly more tumor cells survive chemotherapy and persist through the non-palpable MRD stage in immunodeficient mice. Histological analysis confirmed these findings (**Figure 6B,C**). While well detectable tumor cell islands were observed in the tumor stroma of hematoxylin-eosin (H&E) stained mammary fat pads in both nude and NSG hosts at the MRD stage (**Figure 6B**), the residual tumors in FVB/N mice showed high accumulation of stroma with only a few surviving tumor cells scattered as solitary cells within the tumor stroma, as evidenced by cytokeratin 14 (CK14) immunostaining (**Figure 6C**). Interestingly, during the MRD stage, KB1P tumor cells maintained the expression of the epithelial marker E-cadherin in immunodeficient hosts (**Figure 6B**), whereas most of the rare surviving tumor cells lost E-cadherin expression in FVB/N hosts (**Figure 6C**). As epithelial plasticity and epithelial-to-mesenchymal transition (EMT) contribute to survival of cancer cells after therapy[47], we sought to analyze this further. To monitor the epithelial characteristics of tumor cells, flow cytometry analysis was extended by the epithelial marker EpCAM (**Supplementary Figure S4C,D**). In immunodeficient mice, the vast majority of tumor cells (mCherry+) was also positive for EpCAM throughout the DN, MRD, and REL phases (**Figure 6D**). Importantly, however, in FVB/N mice, while similar results were seen in the DN phase, the few tumor cells obtained at MRD were negative for EpCAM following the TAC(2x,q21) protocol. Treatment with the TAC(2x,q5) protocol also led to significantly diminished EpCAM positivity, albeit with a lower penetrance compared to the TAC(2x,q21) protocol (**Figure 6E**). In REL, the majority of cancer cells reverted to an EpCAM+ epithelial phenotype following both treatment protocols (**Figure 6E**), suggesting a transient loss of epithelial characteristics of a substantial subset of the few surviving tumor cells in immunocompetent hosts.

**Figure 6.**
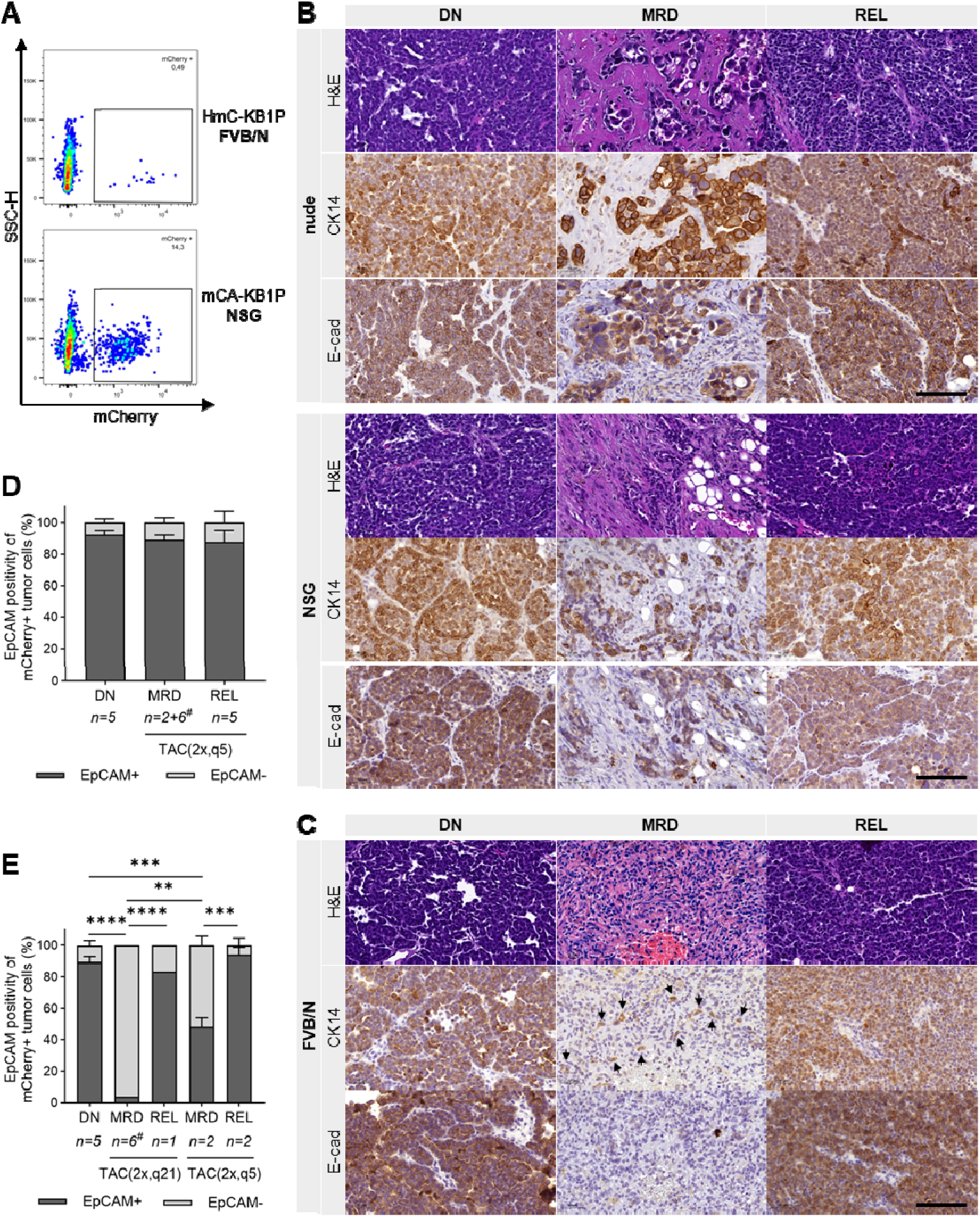
MRD stage following combination chemotherapy is linked to lower selection pressure in immunocompromised mice. **(A)** Quantification of tumor cells (mCherry+) onFACS plots of MRD tumors isolated from FVB/N (top panel, TAC(2x,q21), n=6 pooled) and NSG (bottom panel, TAC(2x,q5), n=6 pooled) hosts. **(B,C)** Representative microscopic images of DN, MRD, and REL **(B)** mCA-KB1P tumor tissues isolated from nude and NSG or **(C)** HmC-KB1P tumor tissues from FVB/N hosts, stained with Hematoxylin-Eosin (H&E) and for Cytokeratin 14 (CK14) and E-cadherin (E-cad) expression. Arrows mark CK14-positive tumor cells. Scale bars, 100 µm. **(D,E)** Percentage ± SEM of EpCAM positivity of mCherry+ tumor cells in DN, MRD and REL of **(D)** mCA-KB1P tumors in immunodeficient and **(E)** HmC-KB1P tumors in FVB/N mice over the TAC treatment protocol. Number of mice used for the analysis is indicated under each bar. Nude and NSG hosts were combined for this analysis (DN: nude n=1, NSG n=4; MRD: nude n=4, NSG n=4; REL: nude n=2, NSG n=3). #: 6 samples were pooled for a single FACS analysis. Two-way ANOVA was used for statistical analysis to compare differences between groups. Asterisks mark significant differences, **** p<0.0001, *** p<0.001, ** p<0.01.

In conclusion, while HmC-KB1P derived tumors in immunocompetent FVB/N mice underwent a transient phenotypical shift during MRD, losing epithelial characteristics, both immunocompromised nude mice (lacking T cells) and immunodeficient NSG mice (with additional defects in B and NK cells and components of the complement system) with mCA-KB1P organoid-derived tumors did not show this transient switch during MRD. This highlights the role of the immune system in shaping adaptive responses and indicates reduced selection pressure on epithelial plasticity in the absence of T cell function.

## 4. Discussion

Model systems, including GEMMs, have been crucial in our understanding of chemotherapy resistance in breast cancer[48, 49]. In particular, the KB1P mouse model has been shown to be a suitable preclinical correlate for treatment resistance in TNBC[21, 23]. Previous studies focusing on the KB1P model have mainly investigated monotherapies, whereas most TNBC patients receive combination therapy based on standard anthracycline regimens[7]. Therefore, this study explored a combination therapy modeling neoadjuvant chemotherapy of TNBC based on the TAC regimen (DTX, DOX, and CP). The approach demonstrated that TAC significantly prolongs the RFS of mice with KB1P organoid derived tumors, mirroring the clinical outcomes observed in patients with TNBC[50].

The small reservoir of cancer cells that persist following treatment, either localized at the primary tumor site, circulating in the bloodstream, or disseminated to distant organs poses a significant challenge to achieving complete remission, as these cells can potentially evade therapy, adapt to the tumor microenvironment, and eventually drive tumor relapse or metastasis[51-53]. The persistent cell population is strongly associated with unfavorable clinical outcomes[54]. A related experimental observation involves DTPs, which serve as an experimental analogue to MRD. These cells may remain undetectable, only to become the source of relapse after long periods of apparent dormancy[55].

To develop a reliable mouse model for evaluating treatment response, we first needed to establish a reproducible response of tumors engrafted from transplanted organoids. While DOX treatment failed to achieve a stable reduction in tumor size, treatment response was improved by Doxil therapy, supporting its use in DOX refractory tumors[24]. Several mice treated with Doxil achieved complete response; however, the timing and onset of MRD remained inconsistent and unpredictable. Therefore, the next step was to implement the TAC regimen with a 21-day treatment cycle, following established protocols[35]. This approach consistently led to complete response, although the timing of the relapse remained unpredictable. Administering the second treatment cycle after 5 days increased the short-term efficacy of the treatment and established a stable MRD phase lasting 30-40 days in 70% of animals, providing a useful model for a synchronized analysis of MRD.

We next sought a highly sensitive method to track DTPs, beginning with the establishment of mCA-KB1P organoids, which allowed detection of as few as 5 cells through AkaBLI. Unexpectedly, mCA-KB1P organoids failed to engraft in FVB/N mice, due to an immunogenic response against AkaLuc. Supporting this hypothesis, AkaBLI also confirmed the elimination of mCA-KB1P organoids in FVB/N hosts. Interestingly, in a subset of nude mice showing no tumors, the engrafted cells persisted in a dormant state for several months (Suppl. Figure 2), indicating that T cells were primarily in charge in FVB/N mice to eliminate KB1P cells expressing AkaLuc. In contrast, all transplantations successfully formed tumors in the severely immunodeficient NSG mice, suggesting that immune cell populations other than T cells, such as B cells, NK cells, and components of the complement system—present in nude but absent in NSG mice—might also contribute to the immune response against the AkaLuc reporter protein by forcing tumor cells to dormancy, independently of T cell function.

The use of NRMI nude and NSG mice required adaptation of the triple-combination TAC chemotherapy, which had not been previously described for these strains. The multifaceted immunodeficiency of NSG mice is known to reduce tolerance towards certain cytostatic drugs[56]. Accordingly, NSG mice in this study tolerated lower doses of DOX and DTX. Despite these limitations, NSG mice demonstrated a stable MRD of reproducible length following TAC treatment, albeit with a shorter median RFS compared to nude mice. Nevertheless, this system allowed the investigation of chemotherapy responses through the DN, MRD, and REL phases and validated the efficacy of the established TAC(2x,q5) treatment protocol in achieving a complete response with stable MRD in the KB1P tumor model across multiple mouse strains, regardless of their immune status.

BLI leverages the catalytic activity of luciferase enzymes on luciferin substrates in the presence of oxygen, magnesium, and ATP to produce light, which is then captured during imaging[57]. Despite several advantages to fluorescence imaging, BLI is constrained by limited imaging depth, challenges in quantifying signals, and reduced spatial resolution [57, 58]. The integration of AkaLumine with the engineered AkaLuc luciferase, referred to as AkaBLI, represents a significant enhancement in imaging sensitivity and depth[29]. Our results showed that AkaBLI has a detection limit of approximately 1,000 cells in mice, which was 10-50 times more sensitive than FLuc BLI, highlighting its potential for *in vivo* MRD detection. However, the sensitivity in organoids *in vitro* was much higher at a detection limit of 5 cells, achieving 100 times greater sensitivity than FLuc BLI. There was also a difference in BLI intensities in the DN compared to the MRD and REL stages. It is possible that the significant metabolic burden imposed by ATP-dependent luciferases such as FLuc and AkaLuc during the bioluminescent reaction, as reported by Yeh et al.[59], might explain stage-dependent differences in BLI signal intensities as a consequence of metabolic adaptation of tumor cells to chemotherapy, but this will require more investigation. Our *in vitro* observation that DTP-stage AkaLuc expressing tumor cells show approximately 4-fold higher BLI signal intensity after DOX treatment than their drug naïve counterparts further supports the hypothesis of drug-induced, tumor cell-intrinsic effects on AkaBLI intensity. This also highlights a limitation, as different phenotypic stages during longitudinal imaging can impact the accuracy of quantification.

This study found that during the MRD phase the tumors were non-palpable for extended periods; however, AkaBLI signals were still detectable, highlighting the sensitivity of this approach. Detailed flow cytometric and immunohistochemical analysis revealed that in immunocompetent hosts, only a very small number of cells were detectable at the MRD stage, and only negligible fraction of those expressed the epithelial markers EpCAM and E-cadherin. This observation suggests that surviving tumor cells are extremely rare and show a transient loss of epithelial characteristics during the MRD phase. In the immunodeficient (NSG/nude) hosts, significantly more tumor cells were detected at the MRD stage, and these cells maintained the expression of the two epithelial markers at similarly high levels as observed in the drug naïve and relapsed stages. Based on these results, we conclude that tumor cells undergo extensive phenotypic changes along the adaptation to the treatment, resulting in the temporary silencing of epithelial markers in the immunocompetent model.

EMT describes a dynamic cellular process wherein epithelial cells lose their defining characteristics, such as apical-basal polarity, intercellular junctions, and E-cadherin expression with concurrent gain of mesenchymal traits that enhance motility[60]. EMT is a significant driver of phenotypic and functional plasticity in cancer, contributing to numerous tumorigenic processes including diminished cell-cell adhesion, increased migratory and invasive capacities, enhanced resistance to anoikis, elevated cancer stem cell-like traits, and distant metastasis[61-63]. Cancer cells undergoing EMT often occupy a spectrum of epithelial and mesenchymal traits, showing inherent plasticity and adaptability[60]. By facilitating phenotypic shifts, EMT enable cancer cells to survive diverse therapeutic pressures, making it a central process in the development of treatment resistance[64, 65]. Cells surviving combination chemotherapy of KB1P tumors appeared to experience a transient loss of the epithelial markers EpCAM and E-cadherin. However, in immunodeficient nude and NSG mice, tumor cells consistently maintained the expression of these genes accross treatment stages, indicating a reduced selection pressure on epithelial plasticity and pointing towards a potential role of the immune system in driving tumor cells to undergo EMT.

Increasing evidence supports the view that DTPs exploit EMT as part of their adaptive response to therapy. For instance, in non-small cell lung cancer (NSCLC), cells treated with the epidermal growth factor receptor (EGFR) inhibitor gefitinib exhibited reduced E-cadherin levels and increased vimentin expression in resistant cells. Similarly, an enriched EMT signature in DTPs was demonstrated for EGFR-mutant lung adenocarcinoma cells[66] and patient-derived melanoma models[67]. Furthermore, DTPs often upregulate EMT-related markers, including ZEB1, Twist, and Slug, following therapy[68-70], which is associated with their survival and adaptability[71]. Similar activation of EMT programs has been observed in HER2-amplified breast cancer cells treated with lapatinib, basal-like breast cancer cells exposed to PI3K/mTOR inhibitors, and paclitaxel-resistant lung or prostate cancer cells tolerating EGFR-TKIs[72-74]. These findings collectively underscore that EMT is intricately linked to the adaptive plasticity of DTPs.

The evolving understanding of TNBC biology has led to the identification of potential therapeutic targets, paving the way for the development of targeted therapies[5]. Nonetheless, challenges persist in identifying predictive biomarkers of response and effectively overcoming therapeutic resistance. This study provides a mouse model system to investigate these aspects, and can also assist in the pre-clinical evaluation of targeted treatments. However, we envisage that this system could be further improved in the future. At present, a major limitation in the field of DTP research is the absence of reliable biomarkers to detect, track, and characterize this rare cell population or to predict their potential to transition from a dormant state into fully re-activated, tumor-forming cells. Additionally, to improve clinical applicability, future studies must also enhance detection sensitivity and establish robust correlations between MRD, DTP prevalence, and key clinical outcomes such as progression-free and overall survival[75].

In conclusion, here we present the development of a mouse model of TNBC that predictably underwent MRD and relapse when undergoing TAC therapy that mimics current neoadjuvant chemotherapy protocols. Similar to the clinical situation, TAC extended RFS in this model system. Enhanced AkaLuc bioluminescence imaging enabled, for the first time, an estimation of the number of surviving tumor cells in the MRD state following intensive chemotherapy. Immunocompetent hosts harbored only a few residual cells, which exhibited a transient loss of epithelial characteristics, whereas immunodeficient hosts had greater numbers of cells that survived during the MRD stage while maintaining an epithelial phenotype. These results suggest that the immune system plays a role in shaping tumor cell plasticity, influencing tumor cell survival following chemotherapy. Importantly, this model system provides a useful method of tracking and investigating rare tumor cells, allowing more detailed investigations into the mechanisms underlying treatment resistance.

## Acknowledgements

KB1P breast cancer organoids, isolated from a spontaneous breast tumor of the *BRCA1* and *p53* deficient GEMM model KB1P, were a kind gift from Sven Rottenberg lab (University of Bern). We thank Johannes Reisecker and Aysan Cerag Yahya for supporting Flow Cytometry analysis and the Preclinical Imaging Laboratory of the Department of Biomedical Imaging and Image-guided Therapy, Medical University of Vienna for helpful instructions on IVIS-assisted BLI. CRediT (Contributor Roles Taxonomy) was used to define contributor roles of the authors. Writing support was provided by Melanie Colegrave of Molecular Cell Research.

## Author contributions

Conceptualization: SMS, GS, AC; methodology: SMS, JDC, MAS, MP, AH, MW, JT, AC; formal analysis and investigation: SMS, JDC, AC. Writing: SMS, GS, AC. Supervision: GS, AC. Funding acquisition: SMS, JT, GS, AC. All authors read and approved the final manuscript.

