## Supplemental File for "Enhanced AkaLuc Bioluminescence Imaging for Longitudinal Intravital Monitoring of Minimal Residual Disease in a Murine Model of Triple-negative Breast Cancer"

[#]equally shared contribution

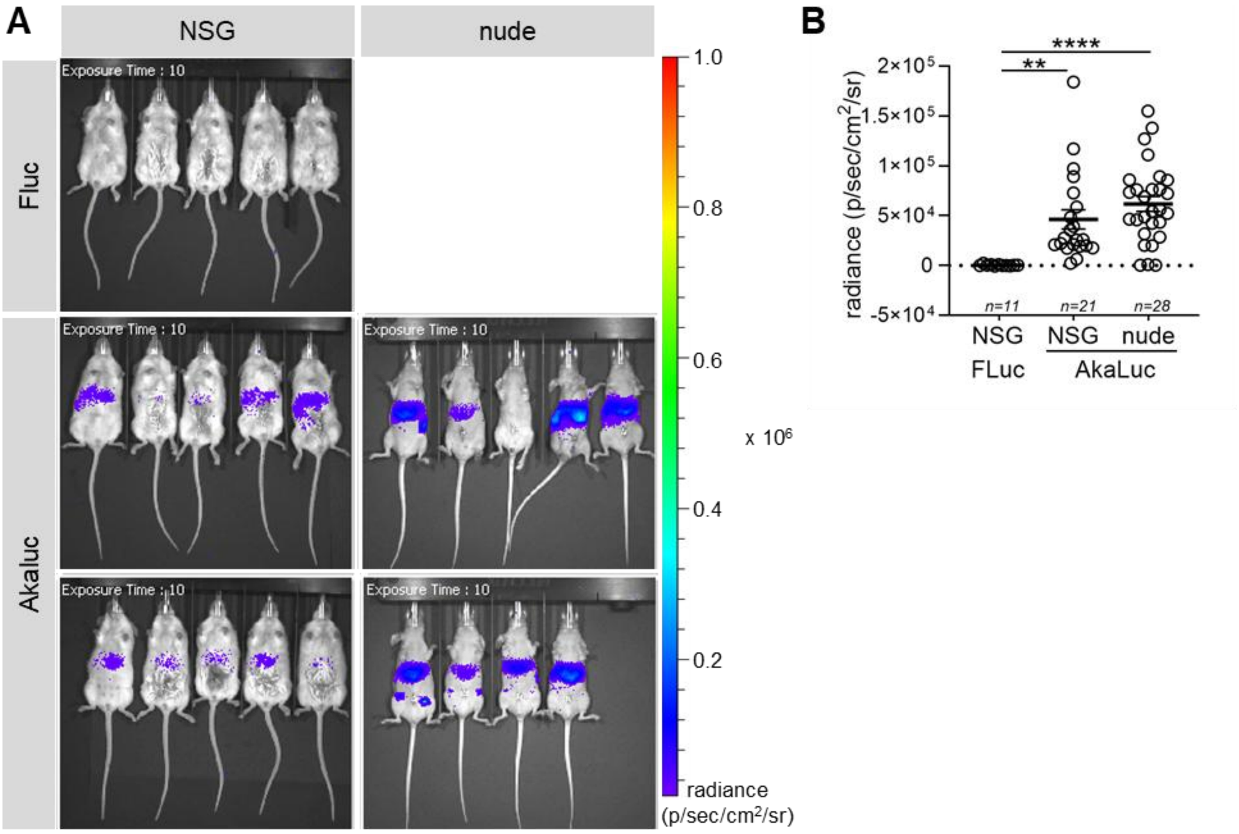


Supplementary Figure S1.

**Supplementary Figure 1 (to Figure 4). The hepatic background of the Akaluc-Akalumine BLI system limits the detection of rare tumor cells in overlapping anatomical regions.**
**(A)** Representative whole-body bioluminescence images of NSG and nude mice from diverse transplantation experiments administered with D-Luciferin or Akalumine substrates.
**(B)** D-Luciferin and Akalumine-derived background bioluminescence of the hepatic area quantified as mean radiance ± SEM in NSG and nude mice.

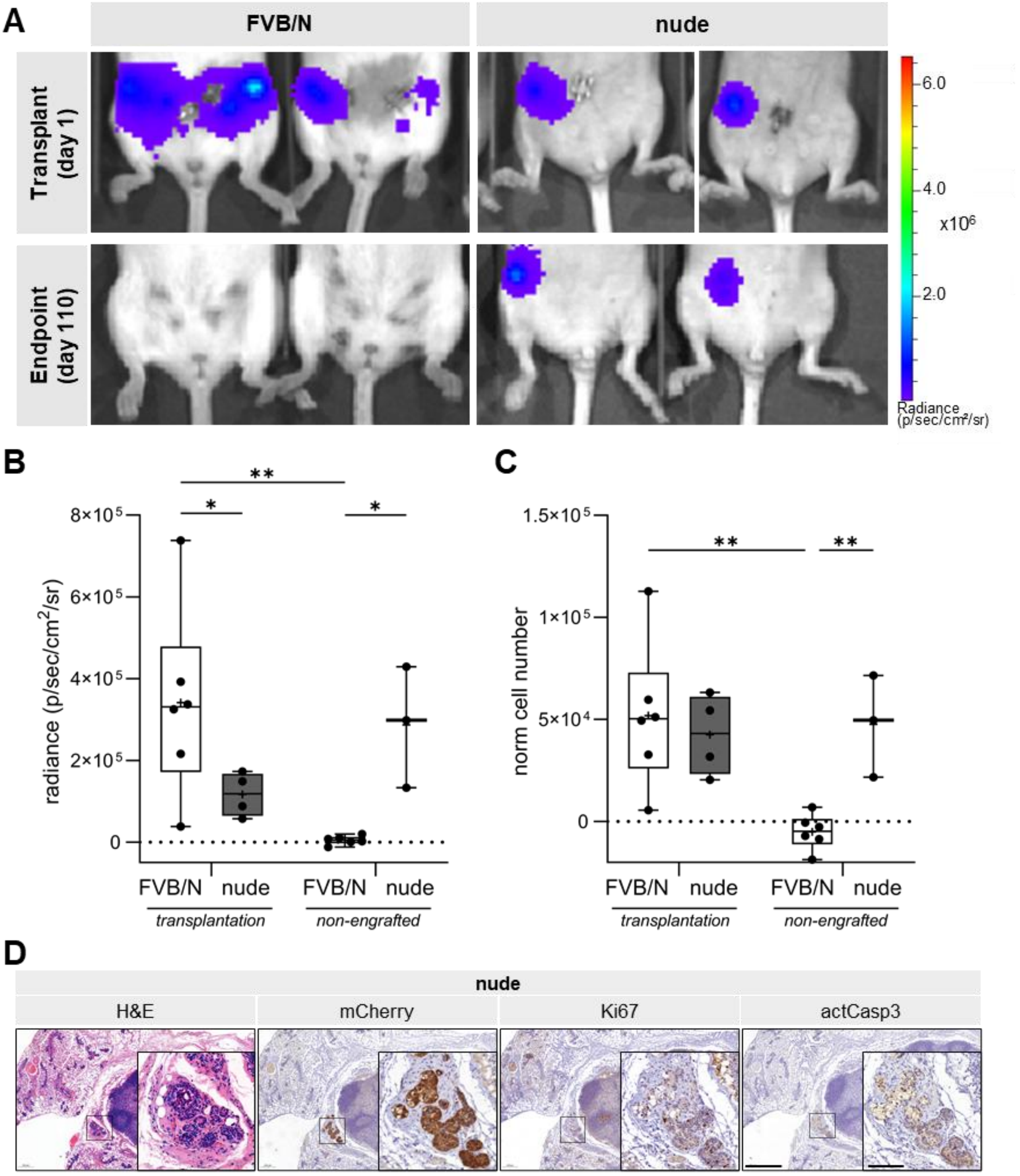


Supplementary Figure S2.

**Supplementary Figure 2 (to Figure 4). Long-term BLI monitoring of mCA-KB1P tumor cells highlights a dormant survival of tumor organoid transplants in nude mice. (A)** BLI images of representative FVB/N and nude mice transplanted with 50,000 mCA-KB1P tumor cells, captured at transplantation (top row) and after 110 days (bottom row). **(B,C)** Quantification on median radiance and corresponding calculated cell number of transplanted and non-engrafted tumor cells based on the dose-curve in Fig.4B. Two-way ANOVA was used for statistical analyses. Significant differences are indicated by asterisks, (** $p<0.01$, * $p<0.05$). **(D)** Representative microscopic images of histological sections of BLI-positive mammary fat pad regions of nude mice without tumor outgrowth 110 days after transplantation of mCA-KB1P tumor cells, stained with Hematoxylin-Eosin (H&E) and for mCherry, Ki67 and activated Caspase3 (actCasp3) expression. Scale bar, 500 µm, for inlet 100 µm.

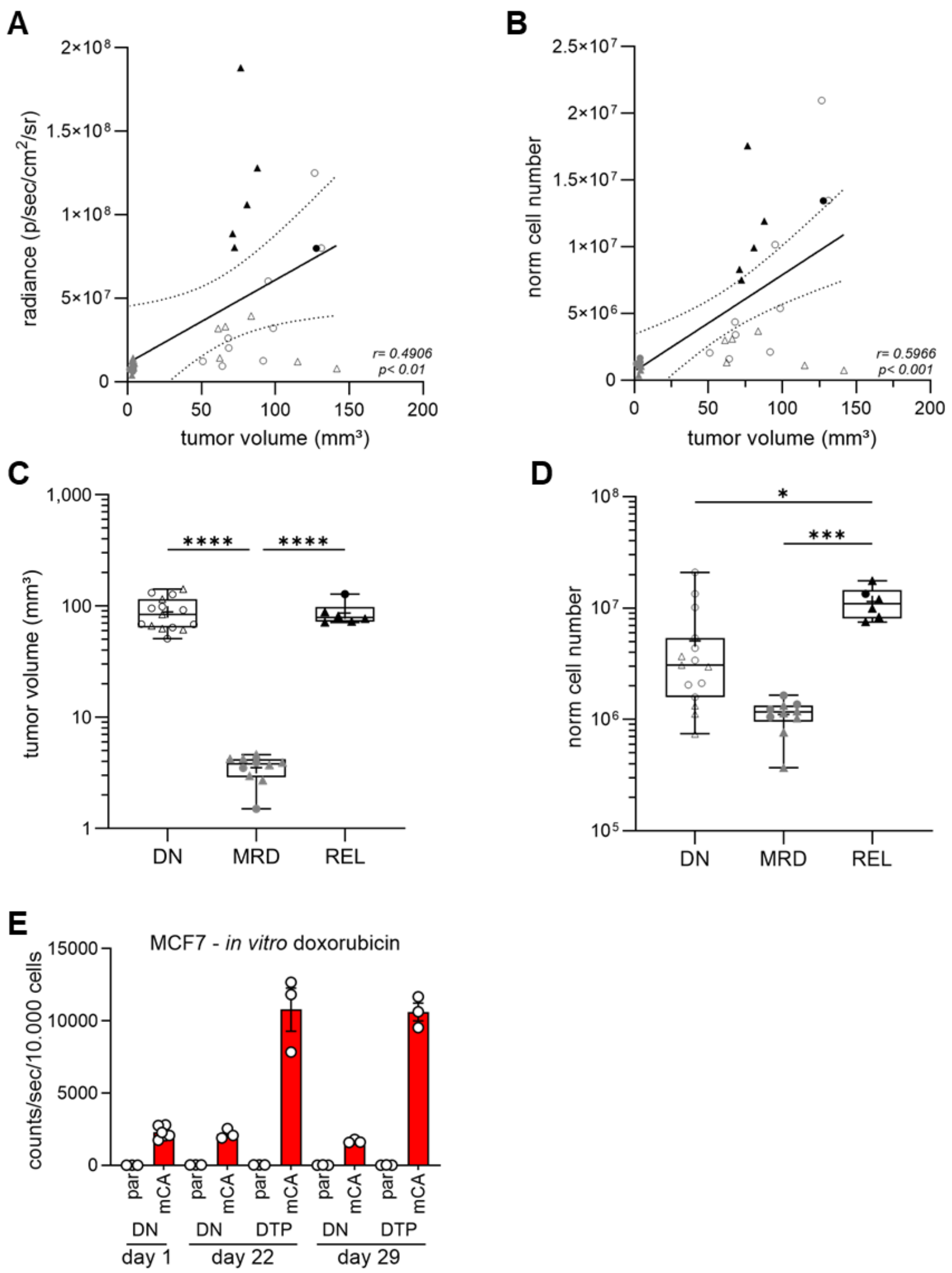


Supplementary Figure S3.

**Supplementary Figure 3 (to Figure 4). (A,B)** Pearson correlation analysis of tumor volume and corresponding **(A)** radiance or **(B)** tumor cell numbers calculated from BLI dose-titration. **(C,D)** Median **(C)** tumor volumes **(D)** and estimated tumor cell numbers calculated from the dose-titration experiments in (A) at DN, MRD and REL stages. **(A-D)** Statistics show combined values for nude and NSG mice. Circles and triangles mark nude and NSG data points, respectively. Empty symbols denote DN, grey symbols mark MRD, and black symbols show REL samples. **(E)** In vitro Akaluc-Akalumine BLI of parental (par) and mCherry-Akaluc overexpressing (mCA) MCF7 cells at different time points of a 2D repopulation assay corresponding to drug naïve (DN) and drug-tolerant persister (DTP) cellular stages quantified as mean counts/sec ± SEM normalized to cell number. **(C,D)** One-way ANOVA with Turkey's multiple comparisons test was used for statistical analyses. Significant differences are indicated by asterisks, (**** p<0.0001).

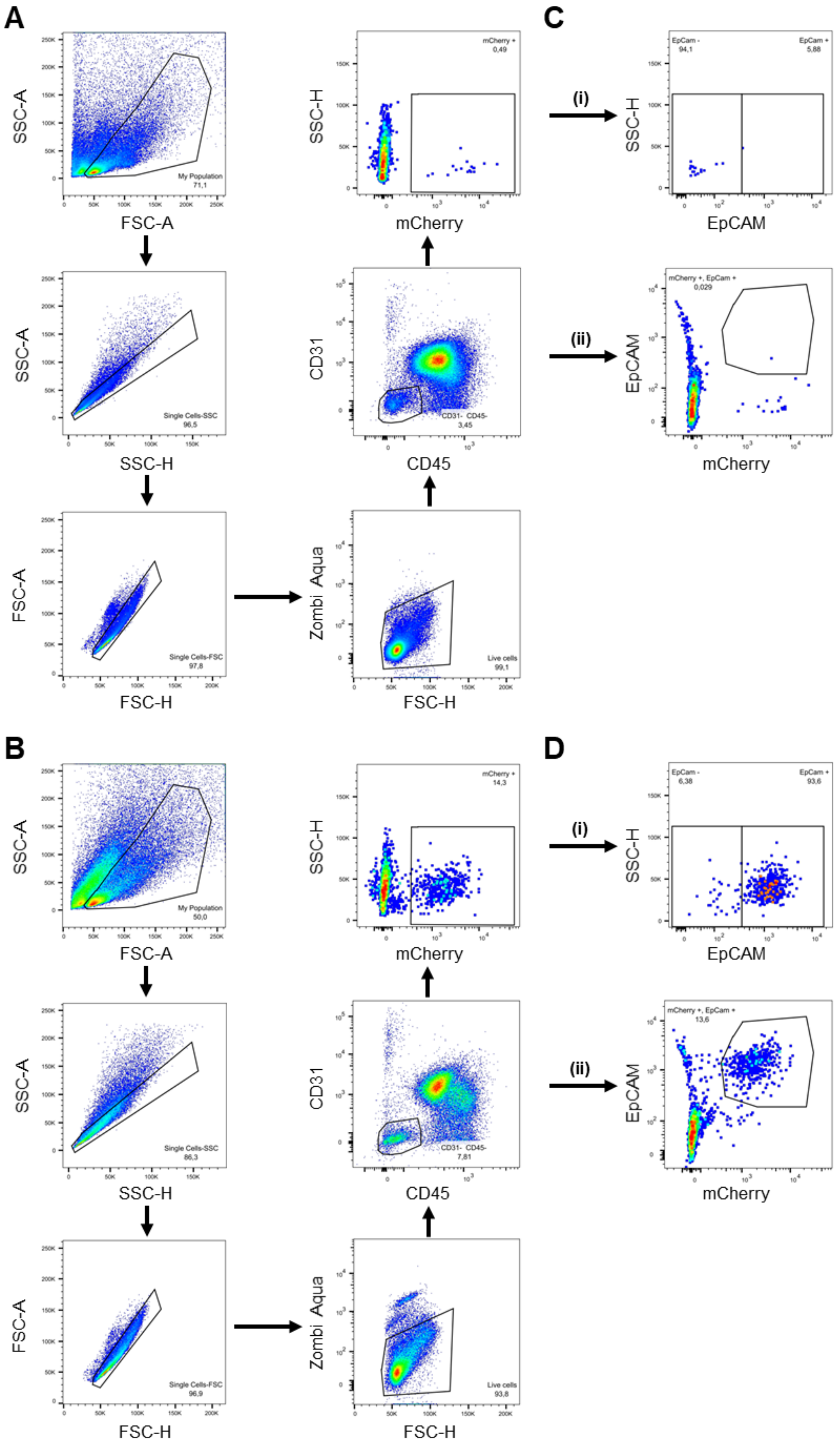


Supplementary Figure S4.

**Supplementary Figure 4 (to Figure 6). Gating strategy for the detection of mCA-KB1P and HmC-KB1P tumor cells and their EpCAM positivity in tumor single cell suspensions.** **(A,B)** Tumor cell detection in **(A)** HmC-KB1P (FVB/N host) and **(B)** mCA-KB1P (NSG host) tumors: events after cell debris exclusion gated for single cell population followed by dead cell exclusion (Zombi Aqua+) and gating the CD45-,CD31- double-negative live cell population for mCherry expression. **(C,D)** EpCAM positivity of **(C)** HmC-KB1P (FVB/N host) and **(D)** mCA-KB1P (NSG host) tumor cells: (i) mCherry positive tumor cells gated for EpCAM expression and percentage of EpCAM- and EpCAM+ tumor cell sub-populations determined; (ii) CD45-,CD31- double-negative live cells gated for EpCAM and mCherry expression to quality control EpCAM immunostaining on mammary epithelial cells of the co-isolated adjacent normal tissue. **(A-D)** show representative MRD stage tumors, same settings were used for DN and REL stages.

**Supplementary Table 1. Median OS and RFS (days) after different chemotherapy protocols in different mouse strains bearing organoid-derived KB1P mammary tumors**

| host | organoid | treatment | | | | | | | | |
|---|---|---|---|---|---|---|---|---|---|---|
| | | untreated | DOX | | Doxil | | TAC(2x,q21) | | TAC(2x,q5) | |
| | | median OS | median OS | median RFS | median OS | median RFS | median OS | median RFS | median OS | median RFS |
| FVB/N | KB1P | 5 | 10 | n.a. | 192 | n.a. | 107.5 | 96 | 45 | 45 |
| | HmC-KB1P | 6.5 | - | - | - | - | 77 | 136 | 52 | 44.5 |
| | mCA-KB1P | no engraftment | | | | | | | | |
| NMRI nude | KB1P | - | - | - | - | - | - | - | - | - |
| | HmC-KB1P | - | - | - | - | - | - | - | - | - |
| | mCA-KB1P | 10 | - | - | - | - | - | - | 59 | 40 |
| NSG | KB1P | - | - | - | - | - | - | - | - | - |
| | HmC-KB1P | - | - | - | - | - | - | - | - | - |
| | mCA-KB1P | 7 | - | - | - | - | - | - | 48 | 33 |

"n.a.": not adequate

"-": experiment not performed
